# Defining the limits of differential transpiration

**DOI:** 10.1101/2025.02.07.637037

**Authors:** Ranjita Sinha, María Ángeles Peláez-Vico, Felix B. Fritschi, Ron Mittler

## Abstract

Differential transpiration is a newly discovered acclimation strategy of annual plants to a combination of water deficit (WD) and heat stress (HS). Under these conditions (*i.e.,* WD+HS), transpiration of vegetative tissues is suppressed in plants such as soybean and tomato, while transpiration of reproductive tissues is not (termed ‘Differential Transpiration’; DT). This newly discovered acclimation process enables the cooling of reproductive organs under conditions of WD+HS, limiting HS-induced damage to plant reproduction. However, at what temperature and WD extremes will this process be active and functional at reducing the internal temperature of reproductive tissues, and at what developmental stages of the plant is it activated, remain unknown. Here, we report that DT occurs at most nodes (leaf developmental stages) of soybean plants subjected to WD+HS, and that it can function under extreme conditions of WD+HS (*i.e.,* 18% of field water capacity and 42°C combined). Our findings reveal that DT is an effective acclimation strategy that protects reproductive processes from extreme conditions of WD+HS, at almost all developmental stages. In addition, our findings suggest that under field conditions DT could also be active in plants subjected to low or mild levels of WD during a heat wave.

## INTRODUCTION

The frequency and intensity of heat waves combined with drought conditions are expected to increase in the coming years due to the intensifying effects of global warming and climate change (Alizadeh et al., 2020; Zandalinas et al., 2021; Lesk et al., 2022; IPCC, 2023). Historically, events of extended droughts combined with heat waves had a devastating impact on crop yield in the US (Mittler, 2006; https://www.ncei.noaa.gov/access/billions/). Compared to conditions of water deficit (WD) or heat stress (HS) applied separately, the combination of WD and HS (*i.e.,* WD+HS), was found to have a greater negative impact on the growth, reproduction, and yield of plants (Rizhsky et al., 2004; Mittler, 2006; Cohen et al., 2021; Sato et al., 2024). This combination was also found to suppress photosynthesis, leaf stomatal conductance, leaf transpiration, and multiple molecular and metabolic pathways of different plants (Mittler and Blumwald, 2010; Zandalinas and Mittler, 2022; Sato et al., 2024). We recently discovered that the combination of WD+HS impacted vegetative and reproductive tissues differently in soybean (*Glycine max*; Sinha et al., 2022, 2023a, 2023b). Thus, while WD+HS caused stomatal closure and suppressed transpiration in vegetative tissues (leaves), in the same plant, these conditions did not cause stomatal closure and suppressed transpiration in reproductive tissues (sepals of flowers and developing pods; Sinha et al., 2022, 2023b). We termed this process ‘Differential Transpiration’ (DT) and identified enhanced rates of abscisic acid degradation and increased stomatal index in developing reproductive tissues as associated with it under conditions of WD+HS. We further found that under conditions of WD+HS the internal temperature of reproductive tissues decreased by about 2-3 °C due to DT (Sinha et al., 2022, 2023b). As the yield of crops such as soybean, corn (*Zea mays*), rice (*Oryza sativa*), and wheat (*Triticum aestivum*) depends on the successful development of reproductive organs, fertilization, embryogenesis and seed filling, and these processes are highly sensitive to HS (Prasad et al., 2008; Djanaguiraman et al., 2018; Johnson et al., 2019; Sze et al., 2021), we hypothesized that DT could protect reproduction in many different crops under conditions of WD+HS (Sinha et al., 2022, 2023b, 2025). Indeed, studies in tomato (*Solanum lycopersicum*) recently revealed that DT is also important for protecting yield under conditions of WD+HS in this crop plant (Bjerring Jensen et al., 2024).

In previous studies we used a defined set of environmental stress conditions to induce WD+HS in soybean, and focused on developing leaves, flowers, and pods from the same developmental stage (node) of soybean plants, to study DT (Sinha et al., 2022, 2023b). Although these studies defined DT and discovered its importance in protecting plant reproduction and yield under conditions of WD+HS, many questions remained unanswered regarding the efficacy of this process under different environmental conditions and at different developmental stages. In the current study we subjected soybean plants to different conditions of WD+HS, as well as studied the occurrence of DT in leaves and flowers developing on different nodes of soybean plants (representing a gradient of development and age differences), to determine the limits under which this acclimation strategy is effective. Our findings reveal that DT is successful in reducing internal flower temperatures under a wide range of WD+HS conditions at almost all developmental stages.

## RESULTS AND DISCUSSION

### Occurrence of differential transpiration at different developmental stages of soybean

To study DT at different developmental stages of leaves and flowers, we grew soybean (*cv Magellan*) plants and subjected them to stress conditions as previously described [Sinha et al., 2022, 2023b; (Control, CT; 55% of field water capacity, 28 day/24 night °C), WD (30% of field water capacity, 28 day/24 night °C), HS (55% of field water capacity, 38 day/30 night °C), and WD+HS (30% of field water capacity, 38 day/30 night °C); Table S1]. Water deficit was initiated when plants reached the R1 stage. Once water-deficit level reached 30% of field capacity, HS (38 day/30 night °C) was applied and two days following the HS application, we measured leaf CO_2_ assimilation, and leaf and whole flower transpiration and stomata conductance, at a range of nodes (leaf developmental) of the growing soybean plants (Figure 1A). As shown in Figure 1B, compared to mature fully expanded leaves (4^th^ and 5^th^ leaves), CO_2_ assimilation was generally higher in younger leaves (1^st^ to 3^rd^ leaves), except in HS (38 day/30 night °C for 2 days). As expected, WD suppressed CO_2_ assimilation, but, interestingly, compared to CT, HS (38 day/30 night °C for 2 days) enhanced CO_2_ assimilation in leaves of all ages (Figure 1B). Compared to WD or HS, however, the combination of WD+HS had the most severe impact on CO_2_ assimilation in almost all leaves (Figure 1B). These findings reveal that, while the level of HS used in this experiment (38 day/30 night °C for 2 days) did not have a negative effect on CO_2_ assimilation of soybean, when combined with WD (WD+HS), the levels of CO_2_ assimilation declined to rates that were mostly lower than CT or WD alone (Figure 1B). This impact of WD+HS on CO_2_ assimilation underlined the unique effects of stress combination on plant physiology (Rizhsky et al., 2004; Mittler, 2006), demonstrating that the rates of CO_2_ assimilation under the WD+HS combination were not the simple average expected from summing the rates of CO_2_ assimilation under WD and HS, rather, they were much lower (Figure 1B). Measurements of transpiration (Figure 1C) and stomatal conductance (Figure 1D) of the different leaves and whole flowers of different ages (Figure 1A), subjected to CT, WD, HS, and WD+HS, revealed that DT between leaves and flowers occurred at all developmental stages along the different nodes measured. Thus, compared to leaves, whole flowers from all different nodes tested under conditions of WD+HS displayed higher levels of transpiration and had a higher stomatal conductance relative to flowers from CT or WD conditions. In contrast, compared to CT conditions, most leaves tested from plants subjected to WD displayed lower levels of transpiration and stomatal conductance, while all leaves and flowers tested from plants subjected to HS displayed higher levels of transpiration and stomatal conductance (Figure 1C, 1D). As previously reported (Sinha et al., 2022), the overall levels of transpiration from whole flowers were about 10-fold lower than those from leaves (Figure 1C). As most plant transpiration occurs through leaves, we calculated what was the decrease in transpiration of leaves and flowers during WD+HS, compared to transpiration levels from these organs during HS (Figure S1). This analysis revealed that compared to HS, leaf transpiration decreased by 95% while flower transpiration decreased by about 40%, under conditions of severe WD+HS stress (WD4+HS) conditions. This finding is important as it demonstrates that plants can save up to 95% of their water intake by using DT to cool their flowers during WD+HS combination, compared to the amount of water needed to cool their flowers and leaves under conditions of HS. Differential transpiration allows therefore the cooling of reproductive tissues even under conditions of limited water availability.

**Figure 1.**
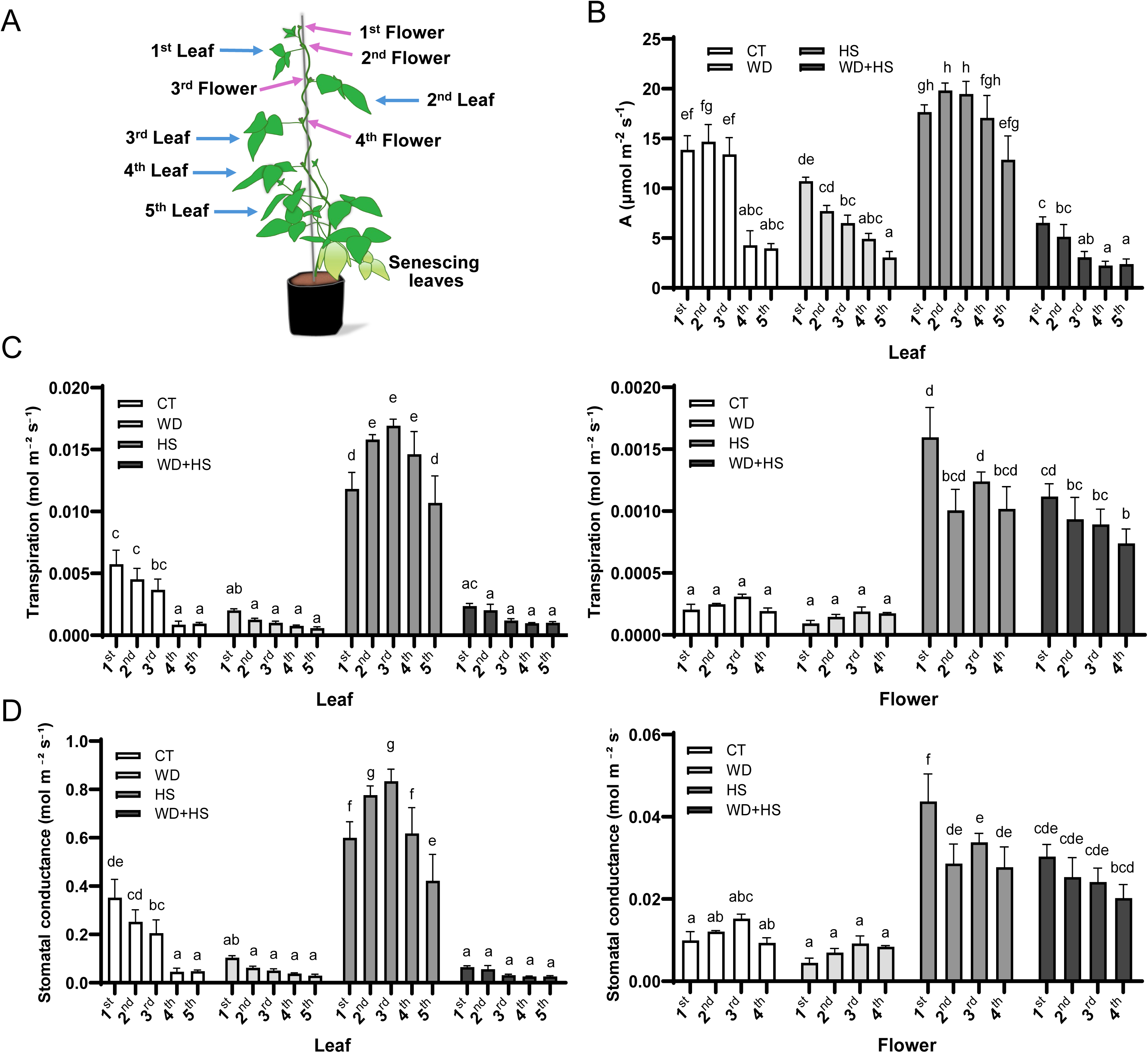
Differential transpiration (DT) at different developmental stages of soybean plants during a combination of water deficit (WD) and heat stress (HS). **A)** A model of a soybean plant showing the different nodes used to study DT at different developmental stages. **B)** Leaf CO_2_ assimilation at the different nodes shown in A, in plants grown under control (CT), WD, HS, and a combination of WD+HS conditions. **C)** Leaf (left) and flower (right) transpiration at the different nodes shown in A in plants grown under CT, WD, HS, or WD+HS conditions. **D)** Same as C but for stomatal conductance. All measurements were repeated in 3 biological repeats, each with 5-7 technical repeats. One-way ANOVA with a Fisher’s LSD post-hoc test was used for significance. Different letters indicate significant at p-value <0.05.

### Establishing an experimental system that subjects soybean plants to different levels of HS, WD, and WD+HS conditions

To study DT under a range of WD and HS conditions, we grew soybean plants until they reached the R1 stage and subjected them to four different WD stress levels (WD1, WD2, WD3, and WD4, corresponding to 40%, 30%, 25%, and 18% of field water capacity, respectively; CT plants were kept at 55% of field water capacity; temperature was kept at 28 day /24 night °C). We then subjected half of the CT and half of the WD1-WD4 plants to a HS of 36 day/30 night °C (HS1, and WD1-4+HS1, respectively) for 2 days. On day 2 of HS1 we measured CO_2_ assimilation of leaves, and stomatal conductance and transpiration of leaves and flowers from node 3 of all plants (Figure 1A). We then subjected all HS1 and WD1-4+HS1 plants to a HS of 38 day/30 night °C (HS2 and WD1-4+HS2, respectively) for 2 days and measured the same physiological parameters described above on day 2. HS2 and WD1-4+HS2 plants were then subjected to a HS of 40 day/32 night °C (HS3 and WD1-4+HS3, respectively) for 2 days and the same physiological parameters measured on day 2 of HS3. The experiment then ended with an additional 2-day stress cycle of 42 day/34 night °C (HS4 and WD1-4+HS4, respectively; Figure 2A; Table S1). The goal of this design (Figures 1A, S2; Table S1) was to simulate a heat wave occurring under different WD conditions. Thus, plants were subjected to increasing levels of HS conditions over 8 days under several different WD levels. As the different HS levels increased in their intensity every 2 days but were transient in each day (temperature linearly increased between 06:00 and 08:00 h, and linearly decreased between 17:00 and 20.00 h; Table S1), this setup also allowed plants to acclimate to the different HS levels (under a set conditions of WD), much like the conditions in the field may allow them to acclimate (Mittler and Blumwald, 2010; Charng et al., 2023; Kambona et al., 2023).

**Figure 2.**
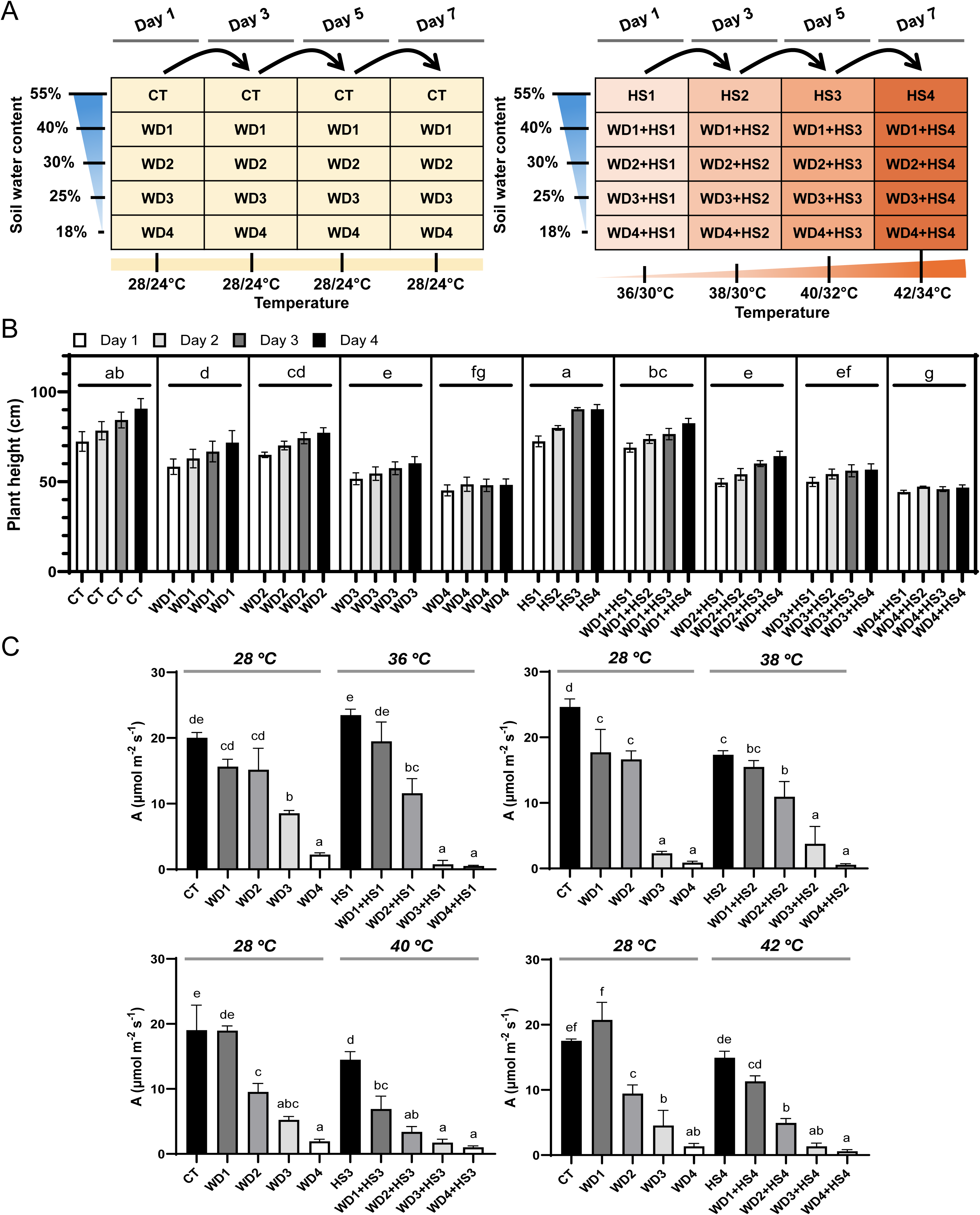
The experimental design used to study differential transpiration (DT) under different levels of heat stress (HS), water deficit (WD), and their combination (WD+HS). **A)** A model showing the different stress conditions and their order. **B)** Changes in plant height during the experiment in plants subjected to the different conditions shown in A. **C)** Leaf CO_2_ assimilation in plants subjected to the different stresses and their combination shown in A. All measurements were repeated in 3 biological repeats, each with 5-7 technical repeats. One-way ANOVA, and Nested-One-way ANOVA, with a Fisher’s LSD post-hoc test, were used for significance in C and B, respectively. Different letters indicate significant at p-value <0.05.

As shown in Figures 2B and S2, while CT plants continued to grow (measured as changes in plant height) during the experiment, the growth of plants subjected to WD (especially WD3 and WD4) was suppressed. Heat stress (applied in the absence of WD) did not suppress the growth of plants, while the different combinations of WD+HS had a negative impact on plant growth (Figures 2B, S2). Measurement of leaf CO_2_ assimilation (at node 3) of the different control and stressed plants revealed that WD3 and WD4, especially in combination with the different HS levels, had the highest suppressive effect on CO_2_ assimilation (Figure 2C). In addition, while a HS of 36 day/30 night °C for 2 days enhanced CO_2_ assimilation, over the course of the experiment and with the increasing levels of HS, CO_2_ assimilation mostly decreased (Figure 2C). At the most severe level of WD (WD4), the negative impact of HS (at any level) on CO_2_ assimilation was the highest (Figure 2C), corresponding with the suppressed growth of plants under these conditions (Figures 2B, S2). The different combinations of WD+HS established using this experimental design (Figure 2A) provided therefore different degrees of WD+HS conditions in which we could measure DT and its effects on internal reproductive tissue temperatures.

### Differential transpiration is effective in reducing internal flower temperatures under a wide range of WD+HS conditions

To determine whether DT is activated in soybean plants in response to different degrees of WD+HS conditions, we measured leaf and flower transpiration and stomatal conductance (at node 3; Figure 1A) of the different plants grown and subjected to the CT, WD, HS, and WD+HS conditions described in Figure 2A. As shown in Figure 3, while conditions of WD3+HS1-4 and WD4+HS1-4 suppressed leaf transpiration (Figure 3A) and stomatal conductance (Figure 3B) under all temperatures, under the same conditions flower transpiration (Figure 3A) and stomatal conductance (Figure 3B) were not suppressed. Overall, compared to CT conditions, flower transpiration and stomatal conductance increased with the increased HS conditions under most WD conditions (HS1-4, WD1-4+HS1-4). This finding suggests that DT could be an inducible stress response process and that the more severe the HS conditions become, plants will conduct more DT under WD+HS conditions (Figure 3). In contrast, leaf stomatal conductance and transpiration decreased in a WD dependent manner under the same levels of HS conditions (WD1-4+HS1-4). The differences in the response of flowers (increasing transpiration with increased HS conditions, regardless of WD conditions) and leaves (decreasing transpiration with increasing WD conditions, regardless of HS conditions) under the different WD+HS conditions, suggested therefore that transpiration and stomatal conductance in flowers and leaves are controlled by different genetic and/or physiological programs. The findings that flower transpiration and stomatal conductance were enhanced under most HS and WD+HS conditions further suggested that flower transpiration plays an important role in protecting plant reproductive processes from overheating under a wide range of environmental stress conditions.

**Figure 3.**
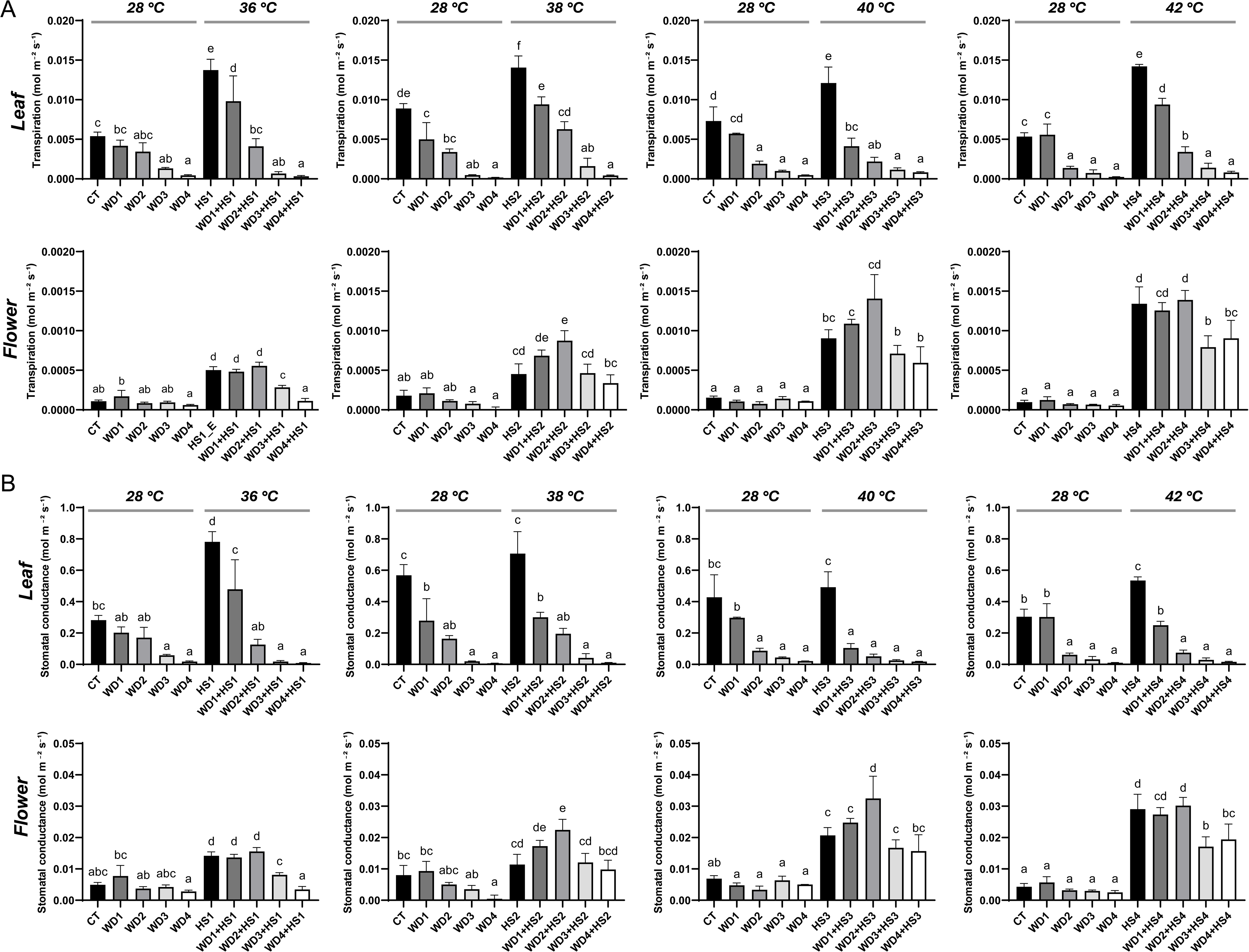
Occurrence of differential transpiration (DT) in soybean plants subjected to a wide range of water deficit (WD) and heat stress (HS) combination conditions. **A)** Leaf (top) and flower (bottom) transpiration of plants subjected to control (CT), WD, HS, and WD+HS as described in Figure 2A. **B)** Same as A, but for stomatal conductance. All measurements were repeated in 3 biological repeats, each with 5-7 technical repeats. One-way ANOVA with a Fisher’s LSD post-hoc test was used for significance. Different letters indicate significant at p-value <0.05.

To determine whether the activation of DT is associated with decreased inner flower temperature (compared to external air temperature), under the different stress conditions, we measured leaf surface temperatures (Figure 4) as well as flower inner (Inner, I) and surrounding air (External, E; at the a distance of about 0.5 to 1 cm from the flower; Figure S3) temperatures (Figure 5), for all conditions shown in Figures 2 and 3. As shown in Figure 4, leaf surface temperature increased with the increased severity of WD and WD+HS conditions. In contrast, flower inner temperature did not (Figure 5). Moreover, compared to the air temperature surrounding the flower, the inner flower temperature was lower by about 2 °C under all HS and WD+HS conditions (Figure 5). Similar differences between surrounding air and inner flower temperatures were largely not observed in plants subjected to CT or different WD conditions, except for a few selected time points/conditions (Figure 5). The findings shown in Figure 5 demonstrate therefore that DT consistently reduces flower inner temperature of soybean plants under a wide range of HS and WD+HS conditions. Whether such a decrease (about 2 °C) would be enough to protect yield under extreme heat conditions in the field remains however to be determined.

**Figure 4.**
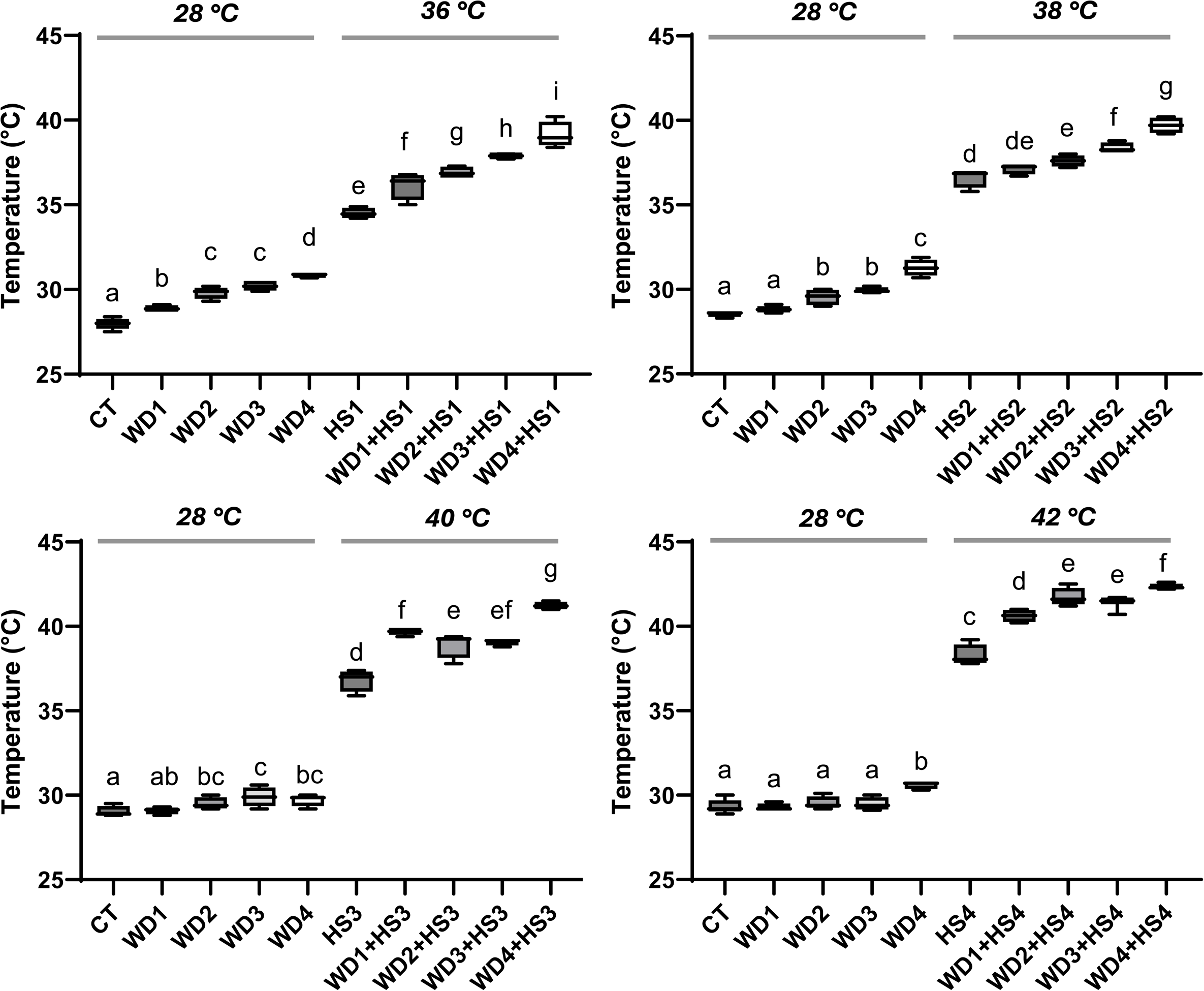
Leaf surface temperature of plants subjected to control (CT), water deficit (WD), heat stress (HS), and WD+HS, as described in Figure 2A. All measurements were repeated in 3 biological repeats, each with 5-7 technical repeats. One-way ANOVA with a Fisher’s LSD post-hoc test was used for significance. Different letters indicate significant at p-value <0.05.

**Figure 5.**
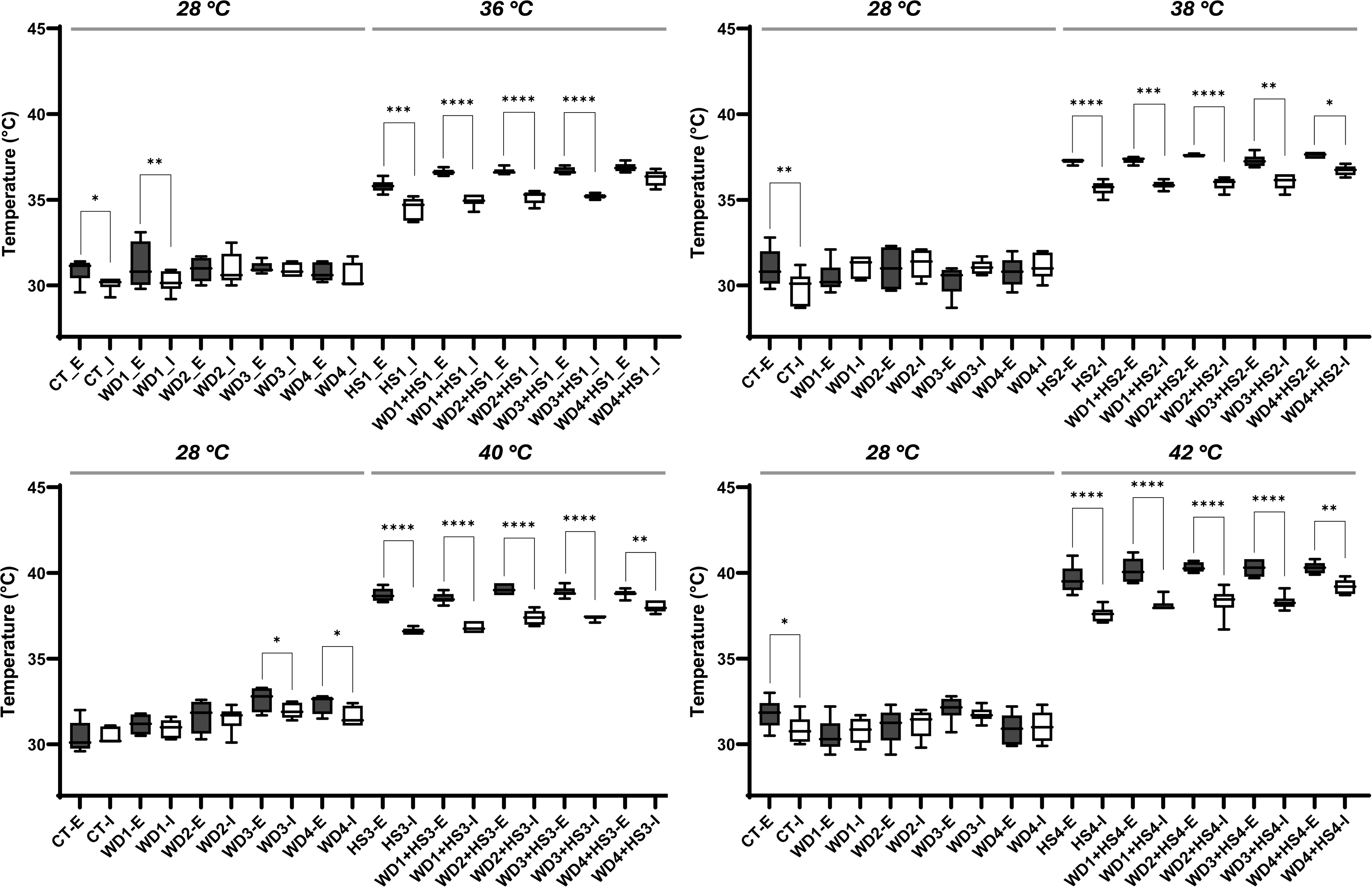
External air and inner flower temperature in soybean plants subjected to a wide range of water deficit (WD) and heat stress (HS) combination conditions. Inner flower temperature (I) and air temperature surrounding flowers (external, E) in plants subjected to CT, WD, HS, and WD+HS as described in Figure 2A. All measurements were repeated in 3 biological repeats, each with 5-7 technical repeats. One-way ANOVA with a Fisher’s LSD post-hoc test was used for significance. Asterisks (*) represent significant difference between external and internal temperature. *, p-value <0.05; **, p-value < 0.005; ***, p-value <0.0005; ****, p-value < 0.0001.

### Summary

The newly discovered acclimation strategy of DT could play a key role in protecting the reproductive processes and yield of our major grain crops from many of the different stresses that accompany global warming, climate change, and increased pollution levels (Zandalinas and Mittler, 2022; Sinha et al., 2022, 2023b, 2025). As many different stresses, such as waterlogging, high ozone levels, and/or pathogen infection, cause overall stomatal closure in plants, the combination of these stresses with HS could pose a similar problem to that experienced by plants subjected to WD+HS, potentially limiting the cooling of reproductive tissues (Sinha et al., 2022, 2025; Zandalinas et al., 2024). It is therefore reasonable to speculate that, in addition to WD+HS, DT could protect reproductive tissues from overheating in plants growing under many other stress combination conditions, and this aspect of DT needs to be addressed in future studies. In the current study we provide evidence that DT is activated at different node positions of soybean plants (Figure 1) and that it is effective in lowering the inner flower temperature of plants under a wide range of HS and WD+HS conditions, including plants subjected to extreme, or low levels of WD under HS conditions (Figures 3-5). This finding suggests that even under ‘natural’ field conditions (*i.e.,* in the absence of severe WD) when plants are routinely subjected to cycles of low or mild levels of WD combined with HS, DT could protect their reproductive tissues from overheating. Additional studies are needed to reveal whether DT is active in protecting developing flowers and pods of different crops from overheating under field conditions, as well as under extreme heat wave conditions (*e.g.,* temperatures in excess of 42-45 °C).

## MATERIALS AND METHODS

### Plant growth and stress treatment

Soybean (*Glycine max*, cv *Magellan*) seeds were inoculated with *Bradyrhizobium japonicum* inoculum (N-DURE, Verdesian Life Sciences, NC, USA) and germinated in Promix BX (Premier Tech Horticulture; PA, USA) mixed with 10% perlite (Miracle-Gro, Marysville, OH, USA), under 14-h light/10-h dark, at 1,000 μmol photons m^-2^ s^-1^ light intensity, 28/24 ℃ day/night temperature (linearly ramped from 24 to 28 °C between 6.00-8.00 AM and linearly decreased to 24 °C from 17.00-20.00 PM), and 60% relative humidity, for a week in growth chambers (PGW40, Conviron, Winnipeg, MB, Canada; Sinha et al., 2022). After a week, seedlings were transplanted into pots containing 1 kg of Promix BX mixed with 10% perlite soaked in 1 L of water-fertilizer (Zack’s classic blossom booster 10-30-20; JR peters Inc., USA; Sinha et al., 2022). Plants were then grown for the next 16-18 days (until start of first open flower, R1; Fehr et al., 1971) under the same conditions. Plants were fertilized twice a week (Zack’s classic blossom booster 10-30-20). At R1, WD was initiated by replenishing only 40%, 30%, 25% and 18% of the water lost by transpiration for WD1, WD2, WD3 and WD4 respectively, while plants in the CT and HS treatments were maintained at 55% of field capacity (Table S1). Once the desired WD level was reached, plants were maintained at this respective WD for the rest of the experiment. For HS treatments, half of the CT and WD1-WD4 plants were subjected to 36/30 ℃ day and night temperature by ramping between 6.00-8.00 AM and decreasing it down to 30 °C between 17.00-20.00 PM for 48 h (HS1 and WD1-4+HS1) and all measurements were taken 27 h after the initiation of HS. Following the 48 h of HS1, all HS1 and WD1-4+HS1 plants were subjected to a HS of 38 day/30 night °C (HS2, and WD1-4+HS2) for a new cycle of HS and WD+HS stresses and all measurements were repeated at the same time. This cycle was repeated for two more time as described in Figure 2A and Table S1. Plant height was measured every alternate day as described by (Assefa et al., 2019).

### Gas exchange measurements

Soybean leaf and flower stomatal conductance, CO_2_ assimilation and transpiration, were measured using a LICOR Portable Photosynthesis System (LI-6800, LICOR, Lincoln, NE, USA) between 11.00 AM-1.00 PM as described previously (Sinha et al., 2022). All measurements were repeated in 3 biological repeats, each with 5-7 technical replicates.

### Leaf and flower temperature

Flower internal (Internal, I) and surrounding air (External; E) temperatures were simultaneously measured with two separate microthermocouple probes (T-29X, Physitemp instruments LLC; Clifton, NJ, USA; Figure S3A; Sinha et al., 2022). Flower internal temperature was measured by inserting one of the microthermocouple probes 1-2 mm into a closed soybean flower bud, while the second probe was kept at about 0.5 - 1 cm away from the flower surface for air temperature surrounding the flower temperature (Figure S3A). In addition, to determine whether shading of the external microprobe will alter its readings (like the shading caused by inserting a microprobe into an unopen soybean flower; Figures S3A), we also measure air temperature in shaded and unshaded probes (Figure S3B; no significant difference). Data was recorded using a Multi-Channel Thermocouple Temperature Data Logger (TCTemp X-Series, ThermoWorks LogMaster; UT, USA) between 11.15 AM-11:45 AM. Surface leaf temperature was recorded using a precision infrared thermometer (DX501-RS, Exergen, Watertown, MA, USA) as described by (Zandalinas et al., 2020).

### Statistical analysis

Graphs are presented as average of 3-5 biological repeats with SEM. One-way ANOVA with Fisher’s Least Significant Difference (Fisher’s LSD) post-hoc test was used for significance. Different letters indicate significant at *p*-value <0.05.

## ACKNOWLEDGMENTS

This work was supported by funding from the National Science Foundation (IOS-2414183; IOS-2110017, IOS-2343815) and the Interdisciplinary Plant Group, and University of Missouri, Columbia.

## AUTHOR CONTRIBUTIONS

R.S. and M.A.P.V. performed experiments and analyzed the data. R.S., M.A.P.V., F.B.F., and R.M. designed experiments, analyzed the data, and/or wrote the manuscript. R.M. and F.B.F. provided financial support.

## DATA AVAILABILITY

The data that supports the findings of this study are available in the text, figure, and supplemental material of this article.

---

**Figure S1.**
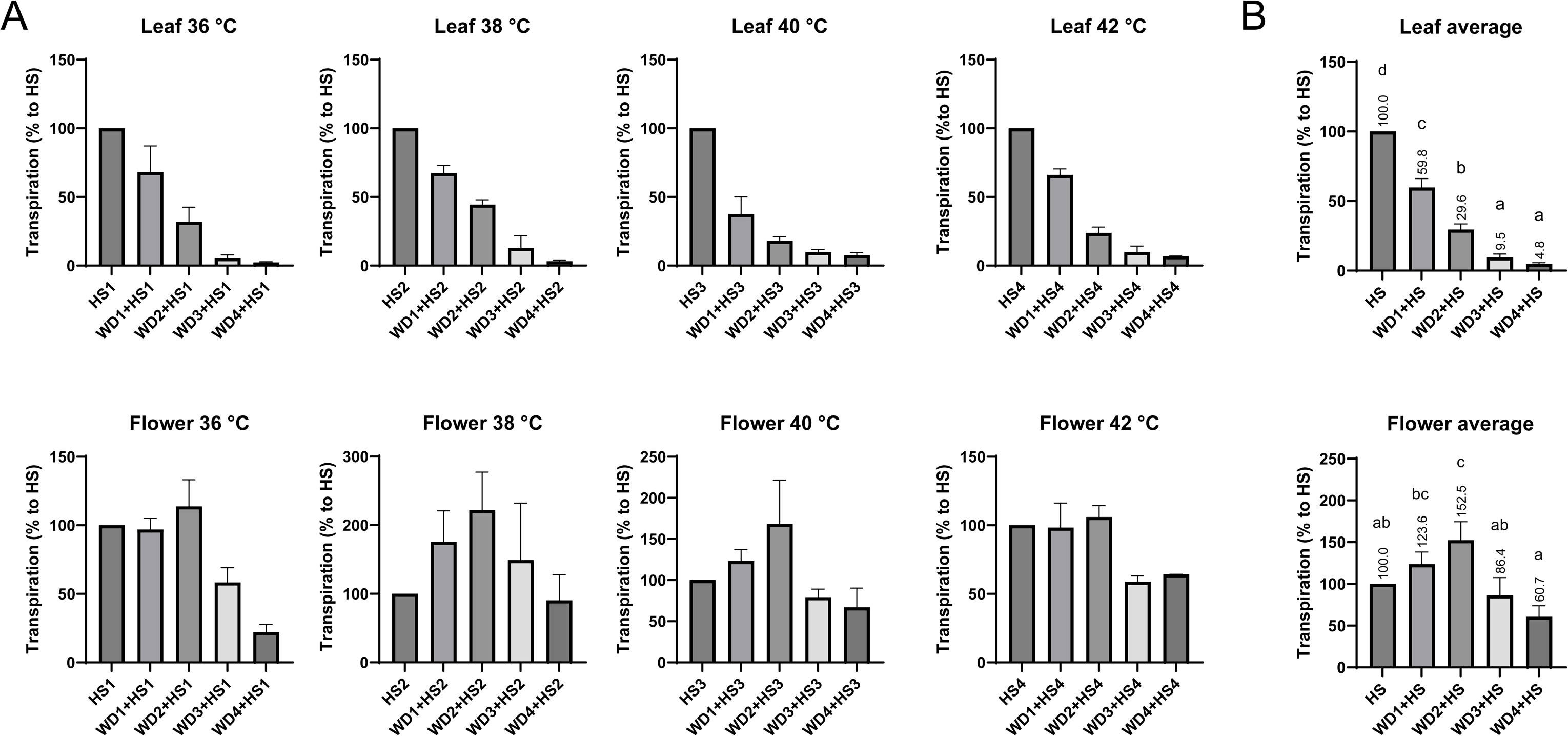
Leaf and flower transpiration in plants subjected to heat stress (HS), water deficit (WD) and a combination of HS+WD, presented as precent of HS transpiration, under each temperature (**A**), or as an average of all temperatures used (**B**). The average of transpiration is given above each bar in (B). This analysis was performed to evaluate the amount of water saved by plants conducting differential transpiration (primarily a result of suppressed leaf transpiration). All measurements were repeated in 3 biological repeats, each with 5-7 technical repeats. One-way ANOVA with a Fisher’s LSD post-hoc test was used for significance. Different letters indicate significant at p-value <0.05.

**Figure S2.**
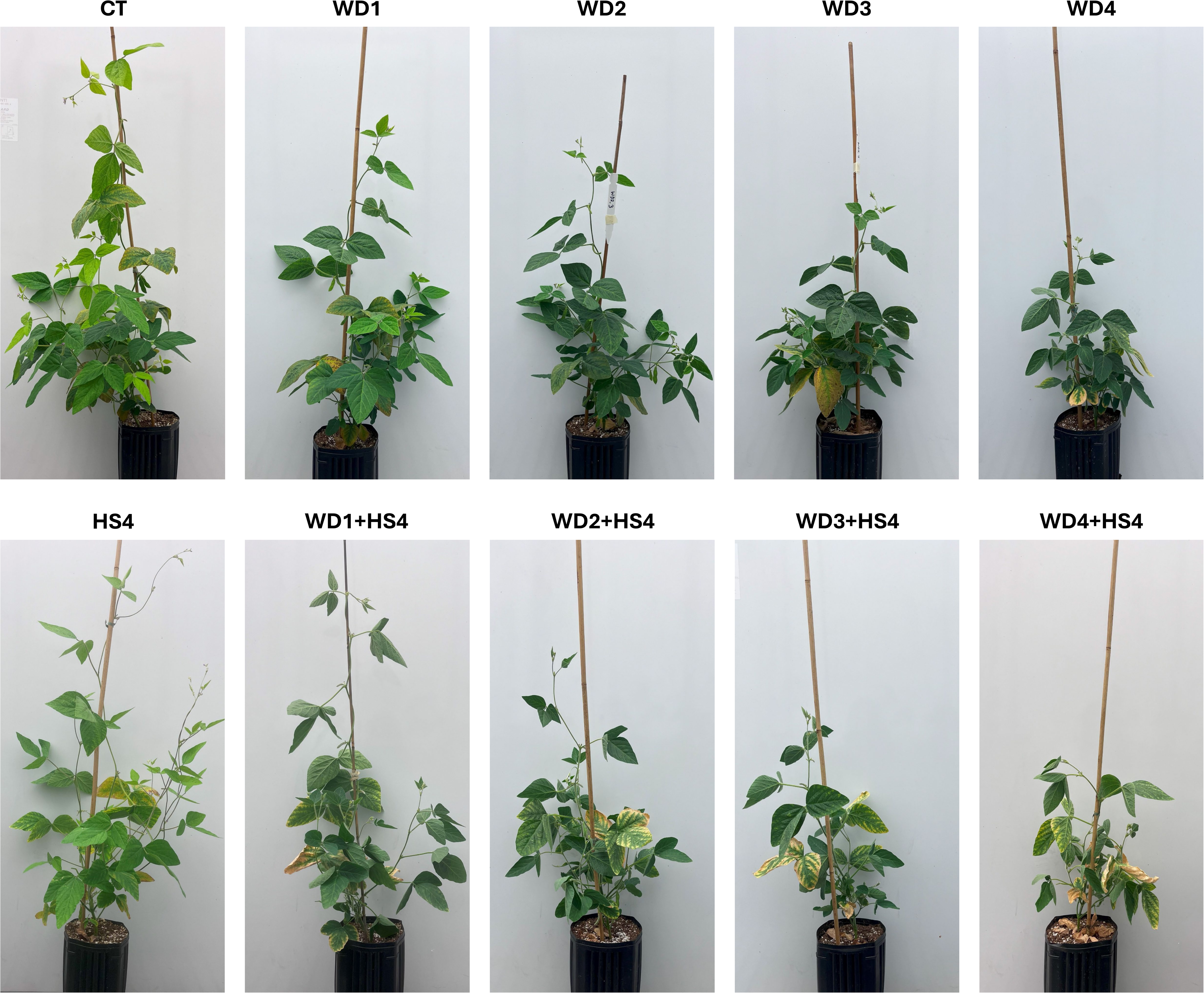
Representative images of soybean plant subjected to the stress conditions described in Figure 2A following the last cycle of HS and WD+HS treatment (HS4 and WD1-4+HS4). Images were taken at noon before WD and WD+HS plants started wilting (typically in the afternoon of each day).

**Figure S3.**
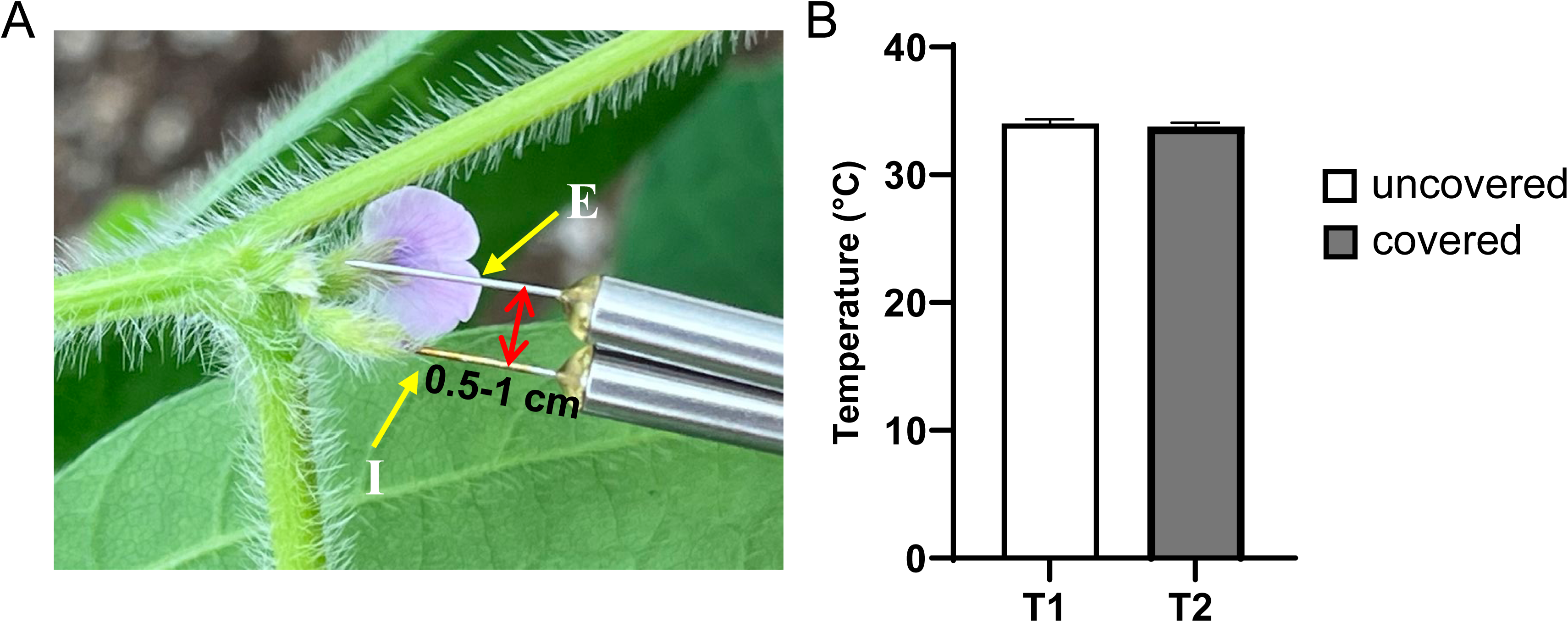
Measurements of flower inner and surrounding air temperatures. **A)** A representative picture of the method used for measuring flower internal (I) and external (E; surrounding air) temperatures using micro-thermocouple probes. **B)** Control analysis of surrounding air (E) temperature with or without shading of the micro-thermocouple probe.

**Table S1:**
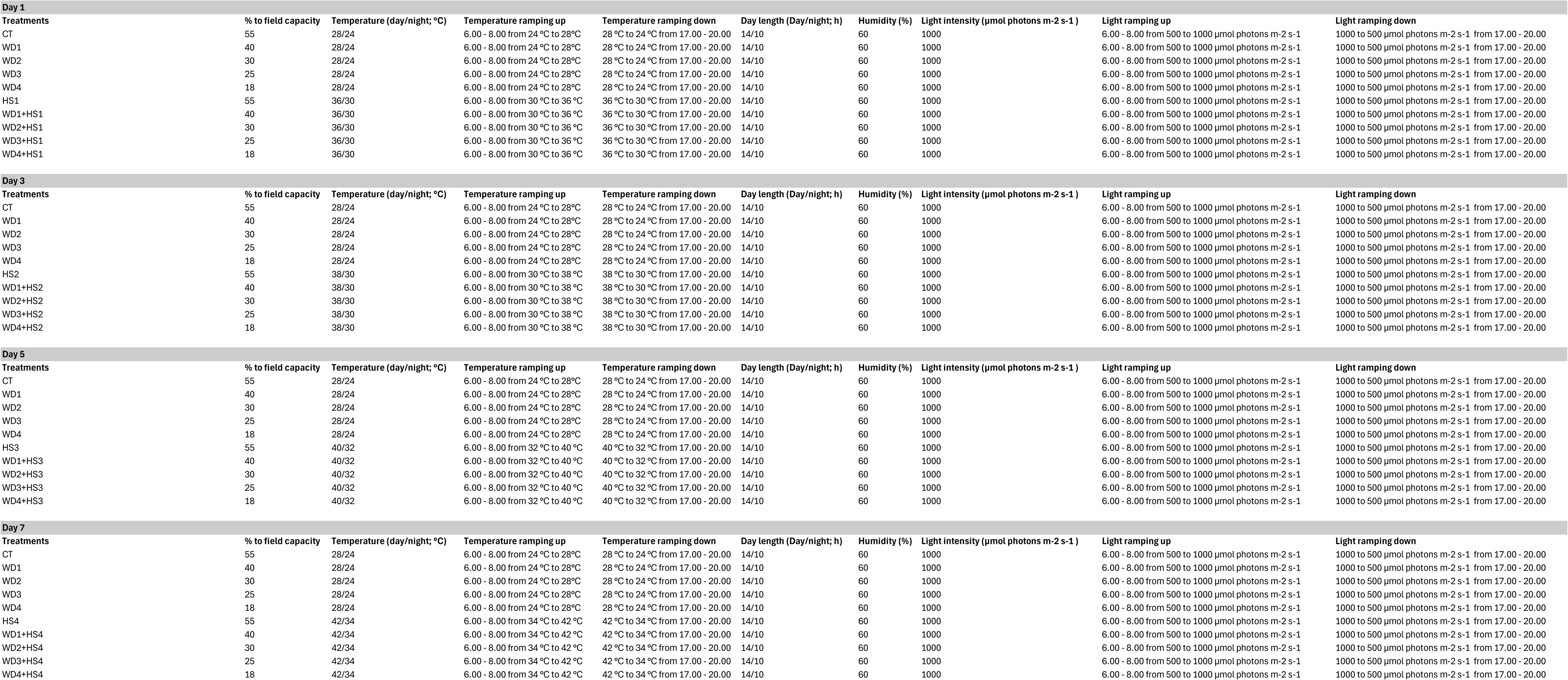
Control and stress treatment conditions used for the experiments shown in Figures 2-5. Control (CT), water deficit (WD), heat stress (HS)

